# Regeneration of lung epithelial cells by Fullerene C_60_ nanoformulation: A possible treatment strategy for acute respiratory distress syndrome (ARDS)

**DOI:** 10.1101/2020.09.08.287680

**Authors:** Nabodita Sinha, Ashwani Kumar Thakur

**Author notes:** Footnotes relating to the title and/or authors should appear here. Electronic Supplementary Information (ESI) available: [details of any supplementary information available should be included here]. See DOI: 10.1039/x0xx00000x.

## Abstract

Acute respiratory distress syndrome (ARDS) involves death of lung epithelial cells. ARDS is a leading reason behind mortality in respiratory infections. Here we show a proof-of-concept that a Fullerene nanoformulation can be used for the regeneration of cells treated with apoptosis-inducing molecules, suggeting its potential for ARDS therapy.

---

Acute respiratory distress syndrome (ARDS) is one of the pathological effects of the SARS-CoV-2 infection which severely damages the lungs. This syndrome also occurs in lung injury, trauma, viral and bacterial infections. However, SARS-CoV-2 induced ARDS is reported to be severe compared to pneumonia or sepsis-induced ARDS.^1^

Lung capillary endothelial cells, type I pneumocytes, and alveolar epithelia get damaged in ARDS, leading to dyspnea and hypoxemia.^2^ Neuromuscular blocking agents and steroids are currently used to reduce inflammation and to increase patient-ventilator synchrony for respiration.^3-5^ Inhaled nitric oxide is another strategy to reduce pulmonary vascular resistance and to facilitate better response to the ventilator. However, these approaches show transient improvement in the patients and exhibit no significant reduction in mortality.^6, 7^

While approaches, such as mesenchymal stem cells (MSCs) and liposomal drug delivery systems have shown some promise in endotoxin (LPS) induced lung injury,^8-10^ novel therapeutic approaches for lung repair and improvement of mortality rate in ARDS is an unmet need in the current times of Covid-19 pandemic. Our lab has recently developed a Fullerene nanoformulation by dispersing 200-400 μg/ml Fullerene powder in cell culture media which results in ^~^170-230 nm (F_P170-230_) sized nanoparticles. This nanoformulation accelerates the proliferation of lung epithelial cells (A549), as well as BMSC, HepG2, NIH-3T3, MDCK and SH-SY5Y cell lines. The nanoformulation was also used in repairing cellular scratch and mice skin wound.^11^ In ARDS, the lung epithelial cells generally undergo apoptosis which ultimately results in the loss of functional cells and consequently impairment of lung function. Regeneration of cell can therefore be one of the therapeutic approaches for ARDS. The basic underlying mechanism of cellular death in ARDS can be successfully modeled *in-vitro*. Cellular death can be induced *in-vitro* by hydrogen peroxide which is a known apoptosis-inducing molecule at certain concentrations.^12^ Toxicity-inducing nanoparticles can be another approach for ensuing cell death. Earlier we have prepared a cytotoxic Fullerene nanoformulation in the size range of 20-50 nm (FT20-50) by dispersing Fullerene at a concentration range of 1-10 μg/ml in cell culture media.^11^ Hence we have used 30 nm sized cytotoxic Fullerene nanoformulation (F_T30_) and hydrogen peroxide to model cellular death in ARDS condition and evaluated the proliferation effect of the Fullerene proliferative (F_P170-200_) nanoformulation for regeneration of lung epithelial cells.

## Effect of the proliferative nanoformulation (F_P170_ and F_P200_) on cell death induced by toxic nanoparticles (F_T30_)

Earlier we had prepared two types of Fullerene nanoformulations and tested them on A549, BMSC, HepG2, NIH-3T3, MDCK and SH-SY5Y cell lines. The toxic nanoformulation of size ^~^20-50 nm (F_T20-50_) showed significant cell death while the proliferative nanoformulation of size ^~^170-230 nm (F_P170-230_) showed 3-4-fold cell proliferation within 24 hours.^11^ **(ESI S1 and S2)** Here the regenerative effects of the proliferative nanoformulation (F_P170_ and F_P200_) on the cells already treated with the toxic nanoformulation was checked. This would test the capability of the proliferative Fullerene nanoformulation to increase cellular viability after insult with a toxic treatment. **(Figure 1 a)**

**Figure 1.**
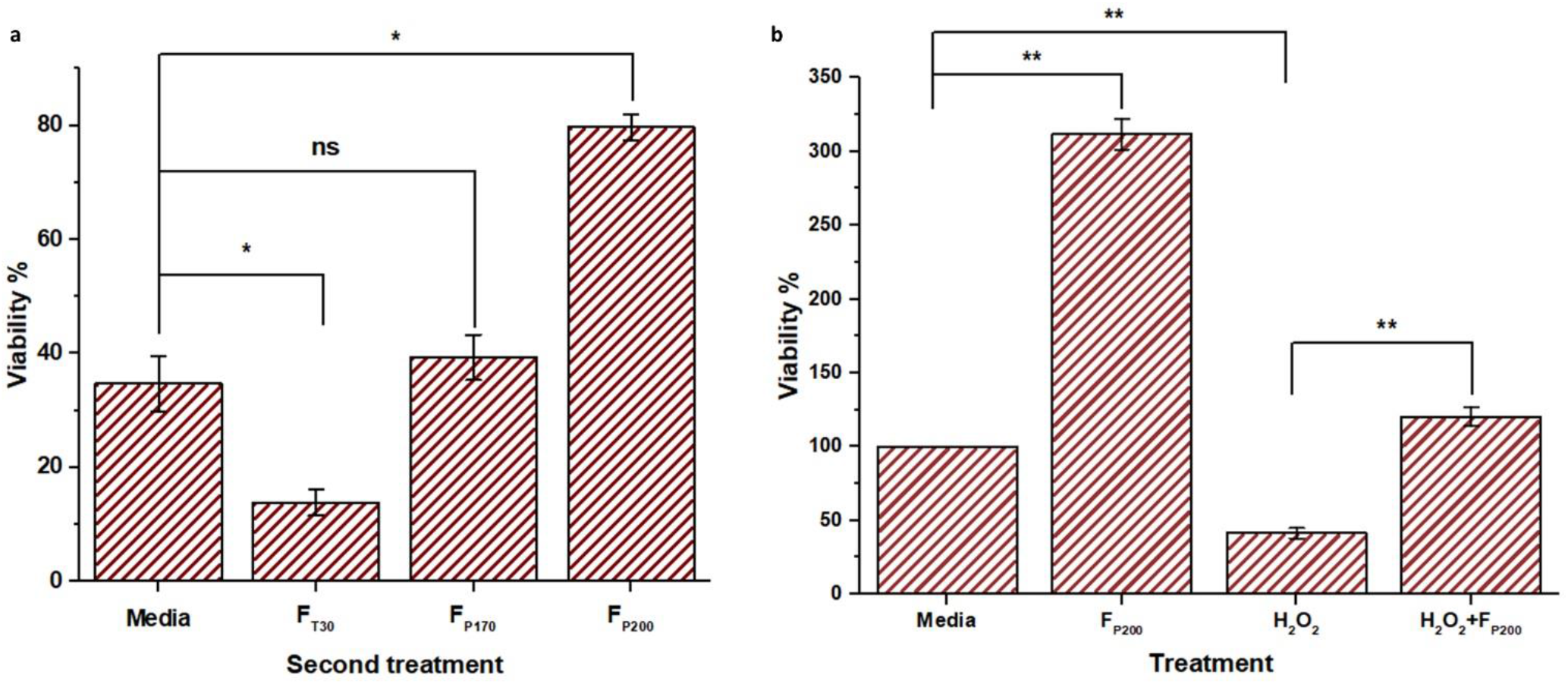
a. The regenerative potential of the proliferative Fullerene nanoformulation on A549 cells after inducing cell death by toxic nanoformulation. First, the cells were treated with 30 nm toxic nanoformulation (F_T30_) for 24 hours. Control cells were untreated for 48 hours and their viability was set at 100% (not shown in the graph). When only media was applied as second treatment, the cell viability became ^~^40% due to initial toxicity treatment. When second treatment was F_T30_, cell viability further reduced to ^~^15%. F_P170_ as second treatment increased cell viability but not significantly compared to control. (t-test; p>0.05) Treatment with 200 nm nanoformulation (F_P200_) showed a significant increase in cell viability (t-test; p<0.05) even after initial toxic treatment (n=3). b. Treatment with F_P200_ nanoformulation alone increased the cell viability up to 300% compared to control. Induction of apoptosis by 0.5 mM hydrogen peroxide decreased the cell viability to 35%. Treatment with both hydrogen peroxide and 200 nm Fullerene nanoformulation (F_P200_) simultaneously increased cell viability up to 125% depicting the potential of F_P200_ to increase cell viability in presence of apoptosis inducing chemical (t-test; p<0.01) (n=3).

First, the lung epithelial cells (A549) were treated with toxic Fullerene nanoparticles of size 30 nm (F_T30_) since in our earlier work these particles have shown significant toxicity on the cell lines tested. **(ESI S4)** After 24 hours, the media was discarded and a second treatment was applied on the F_T30_-treated cells for 24 hours. The viability was then calculated by MTT assay. Viability was normalized to control cells treated with only fresh media for 48 hours (not shown in graph). The entire experiment was repeated thrice (n=3) for statistical analysis. The types of second treatments and the results obtained are as follows: **(Figure 1 a) (ESI S4)**

In the first case (Media bar of Figure 1), the second treatment was only media which was applied after the first treatment of F_T30_. As shown in figure, the viability of the cells is about ^~^40% as compared to 100% viability of the untreated cells. This signifies that after first toxic treatment, treatment with media does not have any significant reparative effect. In the second case (F_T30_) nanoformulation were used as the second treatment. Here the viability was reduced to ^~^15%. Since in the first treatment with F_T30_, viability due to toxic effect reduces to ^~^35%,^11^ the value of 15% viability suggests continuous toxic effect of F_T30_ nanoformulation during second treatment. In the third case, F_P170_ nanoformulation were used as the second treatment. In our earlier work, it was shown that 170 nm particles (F_P170_) can enhance cell proliferation but to a lesser extent than 200 nm nanoformulation (F_P200_).^11^ Here treatment with 170 nm particles (F_P170_) as a second treatment, improved cell viability but not significantly with respect to the case where fresh media was given as the second treatment. (t-test, p>0.05)

In the fourth case, F_P200_ nanoformulation was given as the second treatment. This treatment shows significant increase in cell viability up to ^~^2.3 folds (t-test; p<0.05) even after cell death has been ensued by initial toxic treatment.

Thus here we have shown that even after cellular death is initiated by toxic Fullerene nanoparticles (F_T30_), the cell viability can increase by treatment with proliferative Fullerene nanoformulation suggesting regeneration potential of the proliferative nanoformulation (F_P200_). Next we induced cellular death by a known apoptosis and inflammation-inducing agent hydrogen peroxide and assessed the capability of the proliferative nanoparticles to improve the viability of these damaged cells.

## Induction of apoptosis in cells by hydrogen peroxide and assessment of regenerative capability of proliferative Fullerene nanoformulation (F_P200_)

Hydrogen peroxide at a concentration of 0.5 mM predominantly leads to cell apoptosis.^13^ It is used to induce cell damage and for evaluating the reparative effect of a potential drug candidate. Here we analysed the effect of hydrogen peroxide on the cells with or without the presence of proliferative Fullerene nanoformulation. F_P170_ nanoformulation were not used since it did not show significant proliferative effect in the previous experiment.

The control cells were treated with only media for 24 hours. In the second set, cells were treated with the proliferative Fullerene nanoformulation (F_P200_). When the viability was checked after 24 hours, cells showed an expected 300% increase in cell proliferation as compared to control. In the third set, 0.5 mM hydrogen peroxide was added to the cells. As a result, the viability decreased to ^~^40%. But in the fourth set, when both hydrogen peroxide and proliferative nanoformulation were added simultaneously, the viability improves to ^~^125%. This results in a ^~^3-fold increase in proliferation compared to treatment with only hydrogen peroxide (t-test; p<0.01) **(Figure 1 b).** These results suggest that the proliferative Fullerene nanoformulation can increase the cell viability even after cellular damage by hydrogen peroxide. Similar results were observed in fluorescence microscopy **(Figure 2 a-d) (ESI S5).** The control cells and the F_P200_ **(Figure 2 a and b)** treated cells showed healthy viable cells. 0.5 mM hydrogen peroxide treated cells **(Figure 2 c)** showed a decrease in the number of viable cells. The cells treated with both hydrogen peroxide and F_P200_ simultaneously **(Figure 2 d)** showed a marked increase in the number of viable cells (ImageJ; Analyze Particle Function) compared to hydrogen peroxide alone, signifying the cell proliferative properties of Fullerene nanoformulation. Thus the data obtained suggests regenerative potential of the proliferative Fullerene nanoformulation (F_P200_).

**Figure 2.**
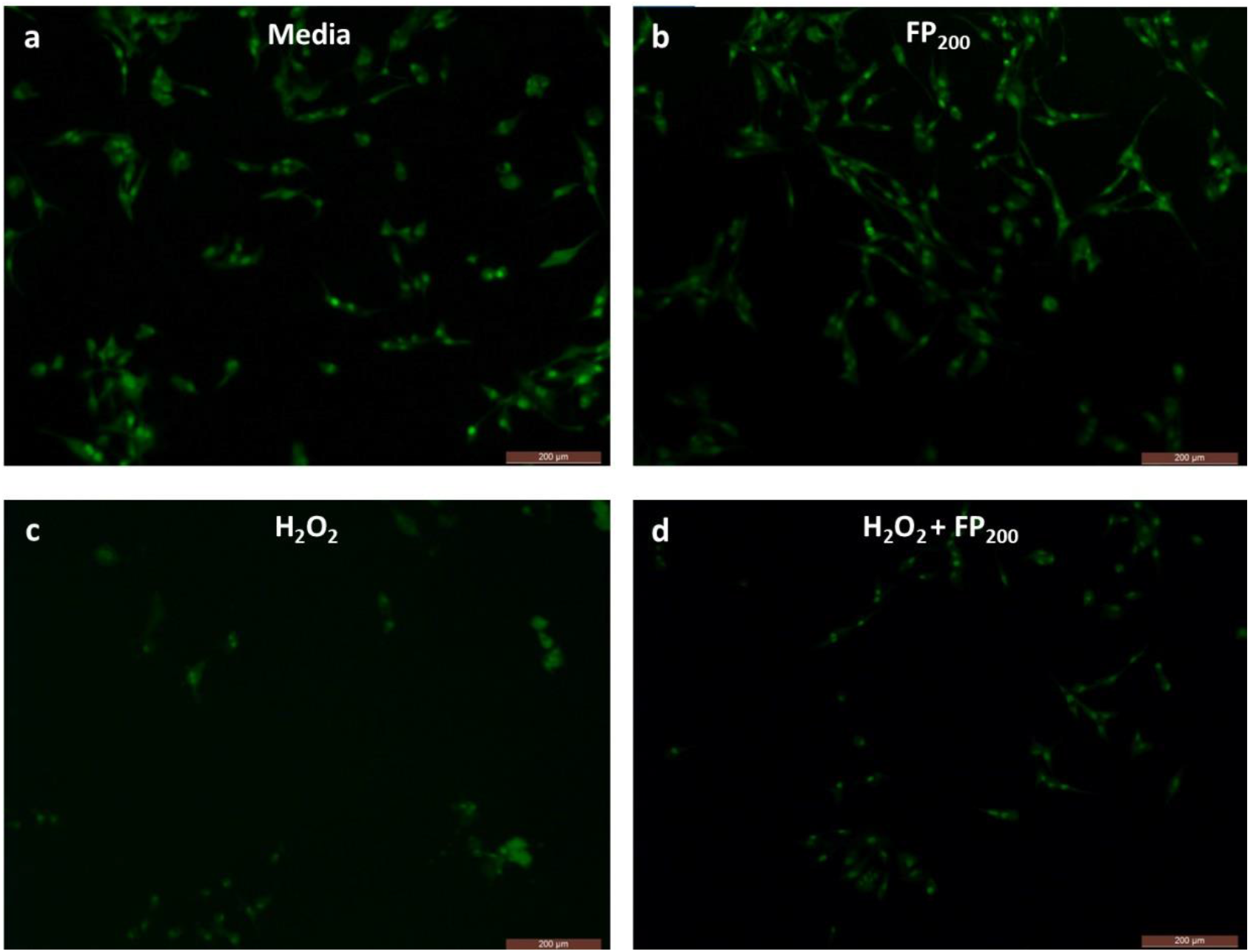
Fluorescence Microscopy analysis of fluorescein diacetate (FDA) stained cells. The viable cells are numerous in both control (a) and F_P200_ treated (b) sample. In 0.5 mM hydrogen peroxide (H_2_O_2_) treated (c) samples, the number of viable cells is very low signifying the cytotoxicity of H_2_O_2_. But cell viability improves when both hydrogen peroxide (H_2_O_2_) and F_P200_ formulations (d) are added simultaneously.

For initial drug development, primary lung epithelial cells, or lung epithelial cell lines can be used to simulate ARDS condition.^14^ Since one of the leading outcomes of ARDS is apoptosis of alveolar epithelial cells,^15^ the end-point of our study was to obtain high cell viability in two experimental models of cellular damage. Here we have shown that the proliferative Fullerene nanoformulation has the potential to increase cell viability after induced cell damage by toxic nanoparticles (F_T30_) or hydrogen peroxide. The improved viability might play a role in mitigating the alveolar epithelial death pathways and might help in restoration of cellular functioning.

In future, rodent models of ARDS would be developed and proliferative nanoformulation would be applied to check lungs repair and mortality of the animals. A thorough toxicological analysis of acute or chronic toxicity of the Fullerene nanoformulation for pulmonary delivery would also be investigated. Encouraging results would further lead to design of an aeroformulation suitable for pulmonary delivery through nebulizer.^15^ The Covid-19 pandemic and resulting ARDS has led to a significant rate of mortality among patients and novel therapeutic approaches in this regard is the need of the hour. The proof-of-concept developed here if successful might prove to be a novel therapeutic approach in reducing ARDS-induced mortality. It would also establish the regenerative potential of Fullerene nanoformulation and would open multiple avenues for exploration of its potential.

## Supporting information

Electronic Supplementary Information

## Acknowledgements

A.K.T gratefully acknowledges financial support from Indian Institute of Technology (IIT) Kanpur, India for this work.

