## Supplementary material for "Regeneration of lung epithelial cells by Fullerene C_60_ nanoformulation: A possible treatment strategy for acute respiratory distress syndrome (ARDS)": Electronic Supplementary Information

**Materials and Methods**

**Instruments Required:** Malvern Zetasizer Nano ZS90 (633 nm laser, scattering angle 90^0^) has been used for DLS measurement analysis. Leica DM 2500 microscope has been used for fluorescence microscopy analysis. For viability assays, Perkin Elmer ELISA Microplate Reader is used for microplate-based tests.

**Detailed methods:**

**S1. Preparation of proliferative nanoformulation (F_P170_ and F_P200_):** In a 15 ml clean glass vial, 2 mg or 4 mg Fullerene powder is weighed and added for preparing 200 µg/ml or 400 µg/ml nanoformulation respectively. 200 µg/ml dispersion produces ˜170 nm sized nanoformulation (F_P170_) and 400 µg/ml dispersion produces ˜200 nm sized nanoformulation (F_P200_). The vial is sterilized by autoclaving at 121°C for 15 minutes. Under aseptic conditions, 10 ml of sterile-filtered DMEM+10% FBS media is added to this vial. The capped vial is stirred for 14 days at 25°C, 700 rpm on a magnetic stirrer.

**S2. Preparation of cytotoxic nanoformulation (F_T30_):** In a 500 ml clean glass bottle, 0.5 mg Fullerene powder is weighed and added for preparing 2.5 µg/ml nanoformulation. This concentration produces ˜30 nm sized nanoformulation (F_T30_). The bottle is sterilized by autoclaving at 121°C for 15 minutes. Under aseptic conditions, 200 ml of sterile-filtered DMEM+10% FBS media is added to this vial. The capped vial is stirred for 14 days at 25°C, 700 rpm on a magnetic stirrer.

**S3. Cell culture:** A549 lung epithelial cell line was used for all the in-vitro experiments. The cells were cultured in DMEM+10% FBS media until cells were 75-80% confluent. For cell growth, 37°C and 10% CO_2_ conditions were maintained. After attaining confluency, the cells were detached from culture flasks using 0.25% Trypsin+0.02% EDTA. For 10 minutes. The detached cells were resuspended in fresh media by centrifuging at 2000 rpm for 1 minute. 90 µL of this cell suspension was mixed with 10 µL of 0.4% of trypan blue. The stained cells were then applied on a hemocytometer and the number of live vs. dead cells were counted. The cell viability is calculated according to the formula below:

**Cell Viability: (Live cell count) / (Dead cell count)**

**Cell Density (no. of cells/mL) = (Average no. of viable cells in each square) ×10000×Dilution factor**

**S4. Effect of the proliferative nanoformulation on cell death induced by toxic nanoparticles:** To 96-well plates, 100 µl cell suspension with a cell density of 20000 cells/ml was added to each well. To these wells, 100 µl of F_T30_ nanoformulation was added and cells were incubated for 24 hours. The used media is then discarded and 100 µl of one of the four types of second treatment was added. The cells were again incubated for 24 hours and the viability was then calculated by MTT assay. For viability normalization, a set of control cells were employed which was treated for 48 hours with only media. For each second treatment type, nine wells are used. The entire set of experiment was repeated thrice (n=3). The treatments are depicted below in the table:

| First treatment | Time | Second Treatment Type | Time |
| --- | --- | --- | --- |
| F_T30_ | 24 Hours | Media | 24 Hours |
|  |  | F_T30_ | 24 Hours |
|  |  | F_P170_ | 24 Hours |
|  |  | F_P200_ | 24 Hours |

**MTT Assay:** 0.5 mg/mL MTT solution (Sigma Aldrich) was prepared in DMEM+10% FBS. 20 μL of MTT was added to each well and the cells were incubated for 2 hours. To the wells, 50 μL of DMSO (Fisher Scientific) was added to lyse the cells. The absorbance was checked at 530 nm using microplate reader. The absorbance values were plotted in Origin Pro 9.1 software. Statistical significance test was performed using t-test (*p<0.05)

**S5. Inducing apoptosis in cells by 0.5 mM hydrogen peroxide and assessment of regenerative capability of proliferative particles**

**S5a. Preparation of 1.5 mM stock hydrogen peroxide solution:** 3% hydrogen peroxide solution was obtained from Sigma Aldrich. To prepare a 50 ml stock solution of 1.5 mM hydrogen peroxide solution, 85.2 µl of 3% hydrogen peroxide was mixed with 49.91 ml of water.

**S5b. Viability analysis:** For this protocol, 100 µl cells having density of 20000 cells/ml were treated according to the four categories:

a. Cells were treated with only 100 µl media
b. Cells were treated with 50µl media+50 µl proliferative nanoformulation (F_P200_)
c. Cells were treated with 50µl media+50 µl of 1.5 mM hydrogen peroxide
d. Cells were treated with 50µl of 1.5 mM hydrogen peroxide +50 µL of proliferative nanoformulation (F_P200_)

For each treatment, 9 wells of 96-well plates were utilized. The whole experiment was repeated on 3 different days (n=3). After 24 hours of treatment, MTT assay was performed as described in Experiment 2 and viability was calculated. Statistical significance test was performed using t-test (**p<0.01).

**S5b.** **Live cell staining**: For fluorescent microscopy analysis, 500 µl of cells (10000 cells/ml) were added on coverslips in 24-well plates and the above treatments were added as follows:

a. Cells were treated with only media
b. Cells were treated with 250µl media+250 µl Proliferative particles (F_P200_)
c. Cells were treated with 250µl media+250 µl of 1.5 mM hydrogen peroxide
d. Cells were treated with 250µl of 1.5 mM hydrogen peroxide +250 µl of Proliferative particles (F_P200_)

After 24 hours, the coverslips were stained with fluorescein diacetate and were analysed with fluorescent microscope. For each treatment, 3 coverslips were used. The whole experiment was repeated thrice (n=3).
